# Unfazed: parent-of-origin detection for large and small *de novo* variants

**DOI:** 10.1101/2021.02.03.429658

**Authors:** Jonathan R. Belyeu, Thomas A. Sasani, Brent S. Pedersen, Aaron R. Quinlan

## Abstract

**Summary:** *Unfazed* is a command-line tool to determine the parental gamete of origin for *de novo* mutations from paired-end Illumina DNA sequencing reads. *Unfazed* uses variant information for a sequenced trio to identify the parental gamete of origin by linking phase-informative inherited variants to *de novo* mutations using read-based phasing. It achieves a high success rate by chaining reads into haplotype groups, thus increasing the search space for informative sites. Unfazed provides a simple command-line interface and scales well to large inputs, determining parent-of-origin for nearly 30,000 *de novo* variants in under 60 hours.

**Availability:** Unfazed is available at https://github.com/ibelyeu/unfazed.

## 1 Introduction

Identifying the origin parent for DNA variants, known as “phasing”, is an important task for understanding molecular mechanisms that generate mutations. Phasing *de novo* mutations can also reveal the effects of parental sex and age on germline mutation rates (Jónsson *et al*., 2017; Sasani *et al*., 2019), and elucidate parental effects on allele-specific expression (Castel *et al*., 2016). Direct phasing tools, unlike statistical inference methods, typically assemble local haplotypes using overlaps between sequencing reads (Martin *et al*., 2016; Edge *et al*., 2017; Hager *et al*., 2020). Although these tools can assign variants to one of two possible haplotypes, they do not directly report the origin parent for those alleles. These tools also are generally applicable only to single-nucleotide variants (SNVs) and small insertion/deletion (INDEL) variants, not structural variants (SVs), which are rearrangements of at least 50 base pairs. Unfazed applies a novel extended read-based phasing method to *de novo* SNV and INDEL mutations identified in family “trios” (mother, father, child), and uses additional non-read-based phasing information from SNVs internal to deletion or duplication SVs. This allows direct prediction of the origin parent for *de novo* variants of all sizes.

## 2 Application

Unfazed identifies the parental gamete of origin for *de novo* mutations via read-based phasing (**Figure 1A**), using individual reads that contain the *de novo* allele and an allele from a phase-informative variant where the origins of the child’s alleles are identified by inheritance. The gamete of origin for the *de novo* allele is inferred by linkage to a phase-informative allele. Recovery of phase information is thus limited by species heterozygosity and the existence of a phase-informative variant near enough to the *de novo* allele to be overlapped by a sequencing read.

**Figure 1.**
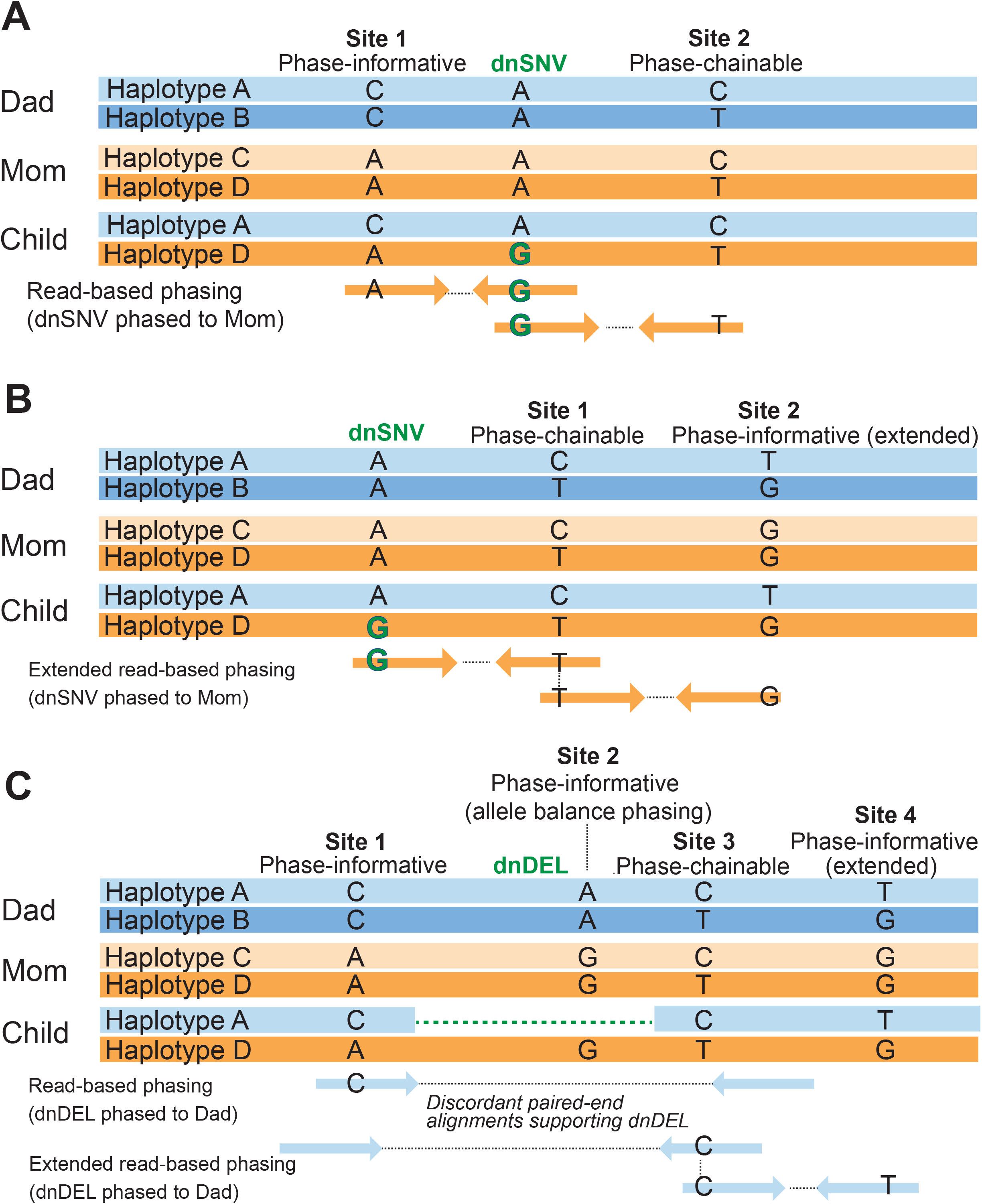
Unfazed identifies the origin parent for variants by extended read-based phasing. **A.** Read-based phasing uses reads overlapping a site of interest and a phase informative site to identify the origin parent. **B.** Extended read-based phasing chains reads to include information from non-overlapped phase-informative sites. **C.** Extended read-based phasing can be applied to SVs by using discordant pairs or split reads.

Unfazed extends the read-based phasing method by including heterozygous loci near the *de novo* allele that are not phase-informative on their own (**Figure 1B**). These loci are “phase-chainable”, meaning they can connect reads overlapping the *de novo* allele with other reads. This increases the potential search distance for informative sites up to kilobases from the de novo site and improves the recovery of phase information by enabling the use of distant phase-informative sites.

Phasing SVs requires specialized logic, as these variants cannot usually be represented by a single Illumina sequencing read. Instead, Unfazed uses logic inspired by SV detection tools (Layer et al., 2014; Belyeu, Chowdhury, et al., 2020) to identify SV evidence in the form of split-reads and discordant read pairs. These reads can then be used to connect the *de novo* SV with phase-informative alleles (**Figure 1C**), allowing the parental gamete of origin to be identified for deletions, duplications, and inversions.

Deletions and duplications, often referred to as copy-number variants (CNVs), change the number of copies of genomic material. This enables another technique for CNV phasing via the allele balance of inherited variants within the CNV. Allele-balance phasing works by finding phase-informative sites inside the CNV (**Figure 1C**) and identifying deviations from the expected allele balance in the offspring. For example, given different homozygous alleles in each parent, the offspring should inherit distinct alleles from each parent. The deletion of one parental copy of a region results in offspring hemizygosity for the other parent’s single-nucleotide alleles in the region. The origin parent for the deletion is therefore the one whose single-nucleotide alleles are lost. Duplications can be phased similarly, using the observation that the duplicated copy includes an extra allele from one parent and increasing the allele balance in favor of the *de novo* mutation’s origin parent.

## 3 Results

Measuring accuracy is challenging for *de novo* variant phasing due to a lack of high-confidence truth sets. However, *de novo* variants from the second generation of a large three-generation pedigree, (Dausset *et al*., 1990) sequenced to 30x coverage and phased using haplotype sharing through all three generations (Sasani *et al*., 2019), contributed a powerful validation set, with large numbers of third-generation offspring ensuring variant transmission. Unfazed reported a parent-of-origin determination for 1,210 out of 4,370 second-generation *de novo* SNVs/INDELs whose origin parent was known from haplotype sharing. The Unfazed prediction was correct in 1,207 cases (99.75%). 7,902 variants were phased by Unfazed from a set of 28,583 unique SNVs/INDEls in both the second and third generations, a phase rate of 27.6% (an increase from 21% with un-extended read-based phasing). Unfazed achieved a phase rate of 40% when applied to a large set of *de novo* SVs (Belyeu, Brand, *et al*., 2020). Command-line example: `unfazed -d denovos.vcf -s snvs.vcf -p pedigree.ped -b bam_directory`

## 4 Discussion

Unfazed is a simple tool for variant phasing with Illumina sequencing reads, with a unique focus on determining the origin parent of *de novo* variants of any size. Unfazed combines ease-of-use and fast runtime with high phase rates for both large and small *de novo* variation. This is accomplished by extending read based phasing to use distant phase-informative sites and leveraging distinct SV read signatures. We anticipate that this tool will prove highly useful to researchers who investigate the rates, patterns, mechanisms, and origins of *de novo* variation.

